# Progressive domain segregation in early embryonic development and underlying correlation to genetic and epigenetic changes

**DOI:** 10.1101/521401

**Authors:** Hui Quan, Hao Tian, Sirui Liu, Yue Xue, Yu Zhang, Wei Xie, Yi Qin Gao

## Abstract

Chromatin undergoes drastic structural organization and epigenetic reprogramming during embryonic development. We present here a consistent view of the chromatin structural change, epigenetic reprogramming and the corresponding sequence dependence in both mouse and human embryo development. The two types of domains, identified earlier as forests and prairies, become spatially segregated during embryonic development, with the exception of zygotic genome activation (ZGA) and implantation, at which notable domain mixing occurs. Structural segregation largely coincides with DNA methylation and gene expression changes. Genes located in mixed prairie domains show proliferation and ectoderm differentiation-related function in ZGA and implantation, respectively. Chromatin of ectoderm shows the weakest and endoderm the strongest domain segregation in germ layers. This chromatin structure difference between different germ layers generally enlarges in further differentiation. The systematic chromatin structure establishment and its sequence-based segregation strongly suggest DNA sequence as a possible driving force for the establishment of chromatin 3D structures which affect profoundly the expression profile. Other possible factors correlated with/influencing chromatin structures, including temperature, germ layers, and cell cycle, were discussed for an understanding of concerted chromatin structure and epigenetic changes in development.

## INTRODUCTION

In mammals, chromatin undergoes drastic organization and epigenetic reprogramming after fertilization(Xu and Xie, 2018; Zheng and Xie, 2019). These processes are essential for gene regulation either globally or specifically by generating chromatin environment that is permissive or repressive to gene expression(Burton and Torres-Padilla, 2014). Chromatin of mouse zygotes and two-cell embryos has obscure high-order structures, existing in markedly relaxed states, which undergoes consolidation of TADs and segregation of chromatin compartments through development(Du et al., 2017; Ke et al., 2017). TAD structure and compartmentalization are also gradually established following human fertilization(Chen et al., 2019). Along with remodeling of 3D chromatin architecture, genome-wide epigenetic reprogramming also takes place during embryonic development(Morgan et al., 2005; Burton and Torres-Padilla, 2010; Dahl et al., 2016; Liu et al., 2016; Zhang et al., 2016; Wang et al., 2018; Xu and Xie, 2018). Global hypomethylation of the genome occurs, and histone modifications globally reset, changing from a non-canonical distribution to canonical one(Smith and Meissner, 2013; Lee et al., 2014; Zheng et al., 2016) during early embryonic development, leading to the specification of the germ layers and cell differentiation.

The progression of the mammalian embryo from fertilization to germ layer formation, concurrent with transcriptional changes and cell fate transitions(Burton and Torres-Padilla, 2014), involves an ordered series of hierarchical lineage determination that ensures the establishment of a blueprint for the whole animal body. One of the most notable transcriptional changes is the zygotic genome activation(ZGA), during which the embryo changes from a state where there is little transcription to another state in which thousands of genes are transcribed(Lu et al., 2016). ZGA is mechanistically coordinated with changes in chromatin state and cell cycle(Jukam et al., 2017). Mammalian ZGA may primarily prepare for the differentiation towards the inner cell mass (ICM) and the trophectoderm (TE), which begins at the morula stage, then the TE can further develop to the extraembryonic tissue necessary for embryo implantation and receiving nutrients. After implantation, the ICM then gives rise to three germ layers (ectoderm, mesoderm and endoderm) through gastrulation, generating founder tissues for subsequent somatic development(Lawson et al., 1991; Arnold and Robertson, 2009).

Thanks to recent developments in low-input chromatin analysis technologies, chromatin structural, epigenetic and transcriptional properties have been roundly explored in the early embryonic development process(Zhang et al., 2018). Taken that chromatin structural properties at the mouse pre-implantation stages have been investigated, analysis on the latest Hi-C data describing chromatin structural properties at post-implantation stages in mouse and embryonic development in human is expected to be able to provide a fairly complete view of the structural change during development. On the other hand, the relationship between global structural changes, epigenetic reprogramming and DNA sequence remains largely unknown and therefore needs to be further investigated.

In our previous study, we analyzed the DNA sequence dependence in the formation of 3D chromatin structures for different cell types(Liu et al., 2018). Based on CpG island (CGI) densities, the genome was divided into alternative forest (high CGI density, F) and prairie (low CGI density, P) domains with average lengths of 1 to 3 million bases (forest and prairie domains are therefore cell type-invariant in one species). CGI forests and prairies effectively separate the linear DNA sequence into domains with distinctly different genetic, epigenetic and structural properties. The segregation of the two domains tends to intensify during embryonic development, cell differentiation and senescence as a result of sequence based thermodynamic stabilization. However, the segregation degree of some somatic cells is less than that of ICM, implying that chromatin structure changes in a non-monotonic way from zygote to differentiated somatic cells. How chromatin conformation gradually changes to establish cell identity during development is thus an interesting open question.

In the present study, we conducted an analysis on global chromosome structural changes during early embryonic development. Two specific stages, ZGA and implantation, during which gene activation and lineage specification occur, respectively, were both found to involve mixing of the two types of genomic domains, with the latter showing a more significant change than the former. The segregation level positively correlates with the proportion of prairie domains reside in compartment B, the larger value of which suggests a more segregated chromatin structure. The DNA methylation distribution in early embryonic development also correlates with the trend of domain segregation in this process, which indicates that chromatins of endoderm and mesoderm are more segregated than that of ectoderm. Moreover, the domain segregation levels of the earliest fate-committed germ layers correlate with those of the differentiated cells. Detailed functions of genes residing in more mixed domains during implantation were then analyzed, based on which we proposed that pluripotent ICM cells may first differentiate to ectoderm-like epiblast via the enhanced interactions of prairies with forests. Finally, we presented a consistent view for the chromatin structural change from birth to senescence, and discussed possible factors influencing global chromatin structure, such as temperature, germ layer, and cell division.

## MATERIALS AND METHOD

### Overall Segregation Ratio

Based on the Hi-C contact matrix, the inter-domain contact ratio between the same domain types was calculated as

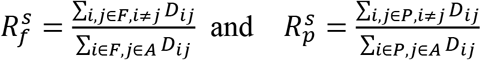

for forests and prairies, respectively. In the above equations, *D*_*ij*_ represents the sum of contacts between the two domains *i* and *j*, which is further divided by the product of domain length of *i* and *j*. *F* is the collection for all forest domains, *P* is the collection for all prairie domains, and *A* is the union of sets *F* and *P*.

The inter-domain contact between different types was calculated similarly as

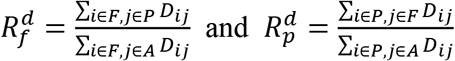

The overall segregation ratio *OR*_*s*_ was then defined as the ratio of inter-domain contacts between the same types and different types:

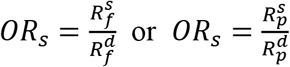

for forests and prairies, respectively. All Hi-C data in this calculation were normalized by ICE normalization(Servant et al., 2015).

### 3D Chromatin Structure Modelling

Detailed information can be seen in our previous work(Xie et al., 2017). Briefly, we first coarse-grained a chromosome as a string of beads. The equilibrium distance between two beads was obtained by converting the contact frequency to the spatial distance. A randomly generated initial structure was then used for further structure optimization using MD (Molecular Dynamics) simulations, until the RMSD (root-mean-square deviation) of the modeled structure became convergent.

### Domain Segregation Ratio Calculation

For each forest/prairie domain *i*, the domain segregation ratio *DR*_*s*_ was defined as the ratio between its inter-domain contacts with the same types and its inter-domain contacts with different types:

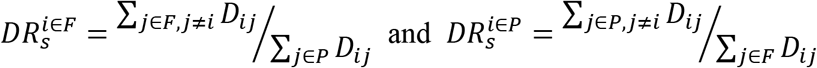

In the above equations, *D*_*ij*_ represents the sum of contacts between the two domains *i* and *j*, *F* is the collection of all forest domains, and *P* is the collection of all prairie domains.

### Distance-dependent Segregation Ratio Calculation

The segregation ratio at distance *d* was calculated as

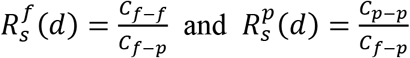

for forests and prairies, respectively. In the above definitions, *C*_*f*−*f*_, *C*_*f*−*p*_ and *C*_*p*−*p*_ are the average Hi-C contact probabilities between forest and forest, forest and prairie, and prairie and prairie, under genomic distance *d*, respectively.

### Open Sea Methylation Difference Index (MDI) between Forest and Prairie

We quantified the methylation difference between adjacent forests and prairies by

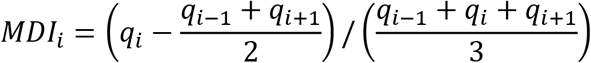

where *q*_*i*_, *q*_*i*−1_ and *q*_*i*+1_ are the average open sea methylation levels for the *i*th domain and its two flanking domains.

### Overall Relative Segregation Ratios

Overall relative segregation ratios (*R*_*or*_) between one tissue and another was identified as

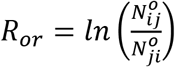

respectively. Here 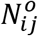 is the number of domains possessing higher *DR*_*s*_ value in tissue *i*, compared to tissue *j*. A positive value of *R*_*or*_ indicates that the chromatin of the former tissue is more segregated than that of the latter. Thus, the parameter *R*_*or*_ is used to generally reflect domain segregation behavior differences between two tissues.

### Significantly More Segregated or More Mixed Domains

To identify forest or prairie domains that are significantly more (or less) segregated in one tissue when compared to another, we first identified threshold values to distinguish more (or less) segregated domains at certain stage. We calculated the logarithm ratio of *DR*_*s*_ values between the two ICM replicates from the same laboratory. Significantly more strongly segregated domains are taken as those whose variation of *DR*_*s*_ values are in the top 2.5 percent tier of all domains, and more mixed domains in the bottom 2.5 percent. The corresponding threshold values of *DR*_*s*_ variation 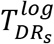 for segregation and mix are then 0.1469 and −0.1469, respectively.

We calculated the logarithm ratio of *DR*_*s*_ between early and late 2-cell embryos for ZGA, ICM and E6.5 epiblast for implantation, respectively. Significantly more strongly segregated domains were identified as those with the logarithm ratio of *DR*_*s*_ higher than 0.1469, and significantly more mixed domains were defined as those with the logarithm ratio of *DR*_*s*_ lower than −0.1469. For convenience, we denoted more strongly segregated and more mixed domains in forest and prairie as *F*_*seg*_, *P*_*seg*_, *F*_*mix*_, and *P*_*mix*_, respectively.

### Compartment Index Calculation

To quantify the degree of compartmentalization, we defined a compartment index (C-index) *I*_*i*_ for 200-kb bin *i* as the logarithm ratio of the average contact between this bin and all A compartments over that between this bin and all B compartments:

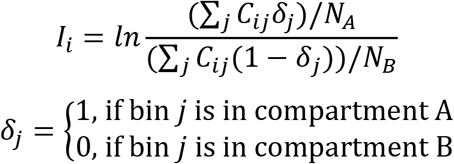

where *C*_*ij*_ is the distance-normalized Hi-C contact probability (normalized by dividing each contact by the average contact probability at that genomic distance) between bins *i* and *j*. *N*_*A*_ and *N*_*B*_ are the total number of bins belonging to compartments A and B, respectively. For each region, a higher value of *I*_*i*_ indicates a more compartment A-like, and a lower one, compartment B-like environment.

### Gene Function Analysis

We analyzed the functional enrichment of genes located in various selected regions by ClusterProfiler and DAVID (https://david.ncifcrf.gov), which yielded similar results. The results were demonstrated via GO terms with p-values. The functional annotation clustering was analyzed using DAVID, and the results were shown via GO terms with enrichment scores.

### Identification of Lineage Specific Genes

To identify lineage specific genes, we used a Shannon-entropy-based method (Xie et al., 2013). Genes with entropy score less than 1.7 were selected as candidates for stage-specific genes. Among them, we selected E6.5 epiblast specific genes satisfying the following conditions: the gene is highly expressed in E6.5 epiblast (FPKM>=1), its relative expression is higher than 1/7, and its expression level in epiblast is higher than that in ICM. These genes were then used for further analysis.

## RESULTS

### Domain Segregation in Early Embryonic Development

Previous studies on chromatin structures during early embryonic development were mainly about changes of structural elements, such as compartment and TADs(Du et al., 2017; Ke et al., 2017). Here we present a structural analysis based on two types of genomic domains to show domain segregation behaviors in different scales. We analyzed chromatin structural changes by calculating overall segregation ratio *OR*_*s*_ (one parameter used to globally reflect chromatin segregation behavior), domain segregation ratio *DR*_*s*_ (which was used to reflect segregation behavior of individual domain), segregation ratio along genomic distances *R*_*s*_(*d*) and F-F (forest-forest) /F-P (forest-prairie) /P-P (prairie-prairie) spatial interaction ratio at each stage.

In our previous study(Liu et al., 2018), based on the division of the mammalian genomes into forest and prairie domains, *OR*_*s*_ was defined as the inter-domain contact ratio between domains of the same types and different types, regarding each domain as one unit (see “Methods”). A higher *OR*_*s*_ for a sample indicates a stronger segregation. Here we calculated *OR*_*s*_ using Hi-C data(Du et al., 2017; Zhang et al., 2018; Chen et al., 2019) of 21 mouse cells and 5 human cells (see Additional file 2: Data sources), which allow us to investigate the chromatin structure changes following the early embryonic development. The calculated *OR*_*s*_ for each stage is given in Figure 1A and 1B. We also constructed the modelled 3D chromatin structures following our previous work(Xie et al., 2017) to show visually the degree of segregation at each stage (see “Methods”).

**Figure 1.**
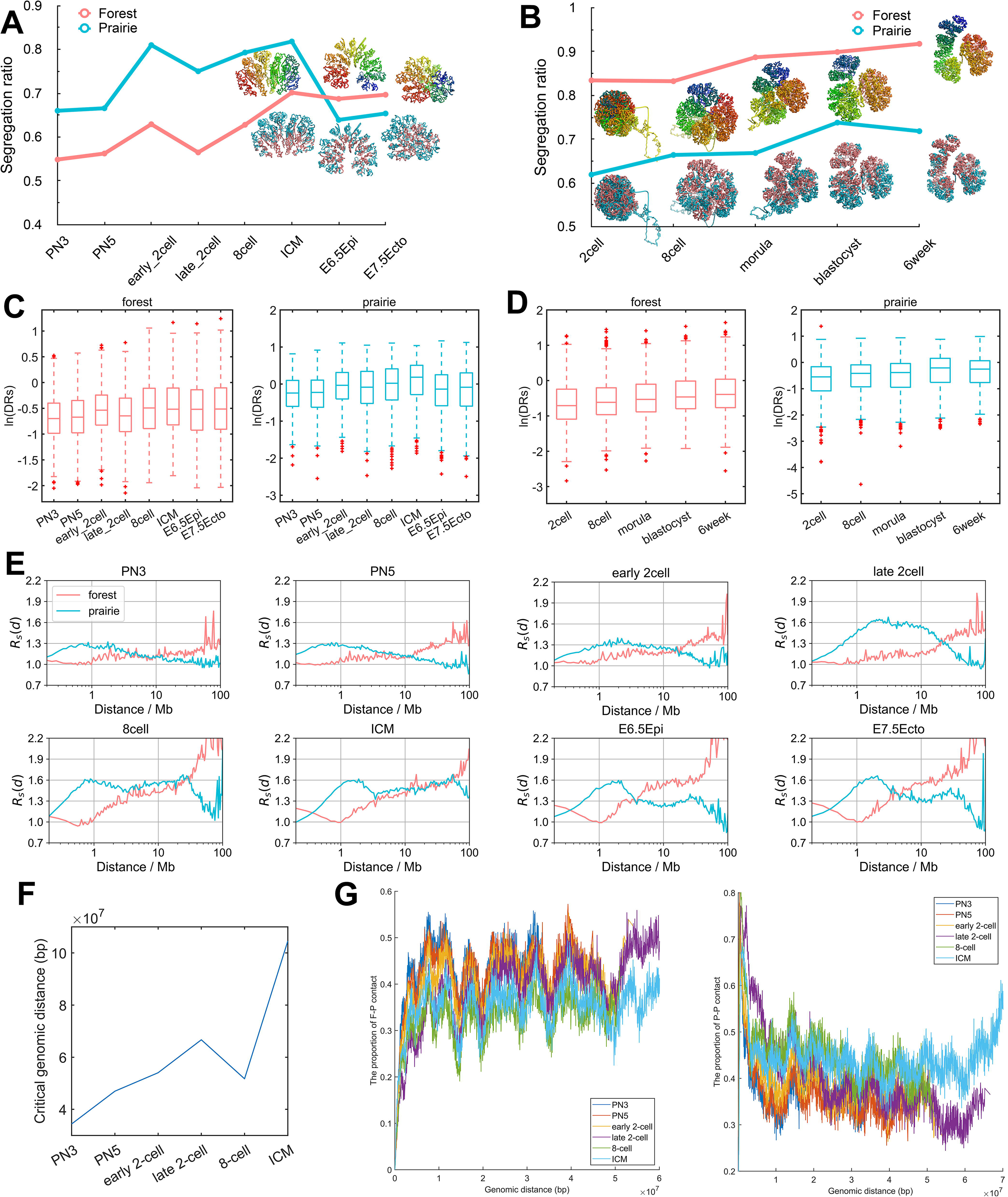
Domain segregation in early embryonic development. **(A, B)** The chromatin overall segregation ratio *OR*_*s*_ of forest and prairie domains at each stage in mouse (A) and human (B) embryonic development. Modelled chromatin structures are also shown at corresponding stages. The upper structures are colored by sequence from blue to red, and the lower structures are colored by forest/prairie domains. **(C, D)** The domain segregation ratio *DR*_*s*_ of forest and prairie domains at each stage in mouse (C) and human (D) embryonic development. **(E)** The distance-dependent segregation ratio *R*_*s*_(*d*) of forest and prairie at each stage in mouse embryonic development. **(F)** The change of critical genomic distance during mouse embryonic development. **(G)** The proportions of F-P and P-P spatial interactions within top 10% contact probabilities under one certain genomic distance at each stage in mouse embryonic development.

As seen from Figure 1A, *OR*_*s*_ increases during the development of early mouse embryonic cells except for two dips. Such a trend suggests the increased segregation of forest and prairie domains from each other, in line with previous observation that more inter-compartment interactions occur at early stages than late stages(Du et al., 2017). The two dips observed along the *OR*_*s*_ curve correspond to the early to late 2-cell, and ICM to E6.5 epiblast transitions, respectively. The decrease during the former transition is seen in both alleles (Figure S1A). For the latter transition, the modelled chromatin structure clearly changes from a domain-separated state to a forest-prairie (F-P) mixed state (Figure 1A). Interestingly, these two processes (that is, early to late 2-cell, and ICM to E6.5 epiblast) exactly coincide with ZGA(Svoboda et al., 2015) and implantation(Bedzhov and Zernicka-Goetz, 2014), respectively. The domain-level F-P chromatin mixing suggested by decrease of *OR*_*s*_ is possibly related to LAD (lamina-associated domains) mixing at ZGA stage as well as germ layer differentiation following implantation, which is analyzed below.

The overall chromatin segregation follows a similar trend in human embryonic development (Figure 1B). *OR*_*s*_ generally increases except for the blastocyst to 6-week transition, accordant with the observed trend of global segregation and domain-level mixing after ICM stage in mouse sample. These changes on domain segregation are also reflected by the 3D structures reconstructed via Hi-C data (Figure 1B). Again, these results suggest that forest and prairie domains tend to segregate from each other before implantation, and domain mixing is observed at the post-implantation stage, showing an increased interaction of the prairie with forest domains.

Besides the overall structure parameter *OR*_*s*_, we also calculated the domain segregation ratio *DR*_*s*_ on each forest/prairie domain during embryonic development (see “Methods”, a higher value indicates the corresponding domain is more segregated), and the variation trend is similar to *OR*_*s*_ (Figures 1C and 1D). Interestingly, the change in *DR*_*s*_ is more significant for the ICM to E6.5 epiblast transition (*p* = 3.7 × 10^−96^ by Welch’s unequal variance test) than the early to late 2-cell transition (*p* = 1.9 × 10^−16^ by Welch’s unequal variance test) for prairie domains, while for forest domains the former one is less pronounced (*p*: 2.4 × 10^−5^ VS 2.1 × 10^−54^, Welch’s unequal variance test).

Further, to investigate chromatin segregation at different genomic distances, we calculated the segregation ratio as a function of genomic distance *R*_*s*_(*d*) (Figures 1E, S1C and S2, see “Methods”). It can be seen from Figure 1E that in early mouse embryo development, prairies are characterized by elevated segregation at genomic distances of several Million bases (Mb). The segregation becomes more pronounced at long distances after 2-cell stage for both forests and prairies, again consistent with an enhancement in domain segregation. Besides, Figure S1B shows a slight decrease of *R*_*s*_(*d*) for forests at the early to late 2-cell stage and a conspicuous decrease of *R*_*s*_(*d*) for prairies at the ICM to E6.5 epiblast stage for mouse, the underlying biological significances behind these phenomena will be discussed below. Enhanced segregation at several Mb scale also occurs during human embryo development (Figure S1C), and an obvious drop of *R*_*s*_(*d*) for prairies at the blastocyst to 6 weeks transition was also observed (Figure S1D).

Finally, the top 10% contact probabilities under genomic distance *d* (that is, the top 10% elements of one diagonal of Hi-C matrix), *TC*_10_(*d*), were extracted, within which the proportions of F-F, F-P and P-P interactions were calculated (notably, the premise of such a calculation is that all elements in the *TC*_10_(*d*) should be non-zero, otherwise the proportions of F-F, F-P and P-P are assigned zero). The genomic distance, after which the proportions of F-F/F-P/P-P remain to be zero, is named the critical genomic distance. During the development of mouse embryo, the critical genomic distance gradually increases (Figure 1F), showing the gradual establishment of long-range chromatin contact. At the same time, F-P ratio descends while P-P ratio increases from PN3 to ICM stages, again hinting that forest and prairie become more spatially segregated along with the development (Figure 1G). These phenomena were also observed in human embryo development (Figure S1E).

### Compartment Changes Relate with Domain Segregation

By means of the Hi-C measurement, chromatin can be partitioned into two spatial compartments (A and B)(Lieberman-Aiden et al., 2009). It was also found that forest domains reside mainly in compartment A and prairie domains mainly in compartment B in different cells, manifesting that DNA with similar sequence properties tends to spatially interact (Liu et al., 2018). Based on these, we calculated the proportion for the two sequential domains (forests and prairies) to be distributed in the two spatial compartments A and B during development.

The corresponding four types of DNA sequences are named as Af, Bf, Ap, and Bp, respectively, with the first letter denoting the compartments and second the forest (f) or prairie (p) domain. One can see from Figure 2A that during mouse development the proportion of prairies in compartment B (Bp) changes in the same trend as the overall segregation ratio, which increases from PN3 to ICM stage, except for the early to late 2-cell stage, then decreases from ICM to E6.5 epiblast, and slightly increases when E6.5 epiblast develops to E7.5 ectoderm. The ratio of Ap changes in a direction opposite to that of Bp. Similar phenomenon was also observed in human embryos, where the changes of *OR*_*s*_ are in accordance with the size of Bp (Figure S3A). Recent study also found that attractions between heterochromatic regions are crucial for compartmentalization and domain segregation of the active and inactive genome(Falk et al., 2019). Similar to it, we found here that during embryo development, the aggregation behavior of prairie domains (low CpG density region) strengthens and more and more prairie regions belong to compartment B.

**Figure 2.**
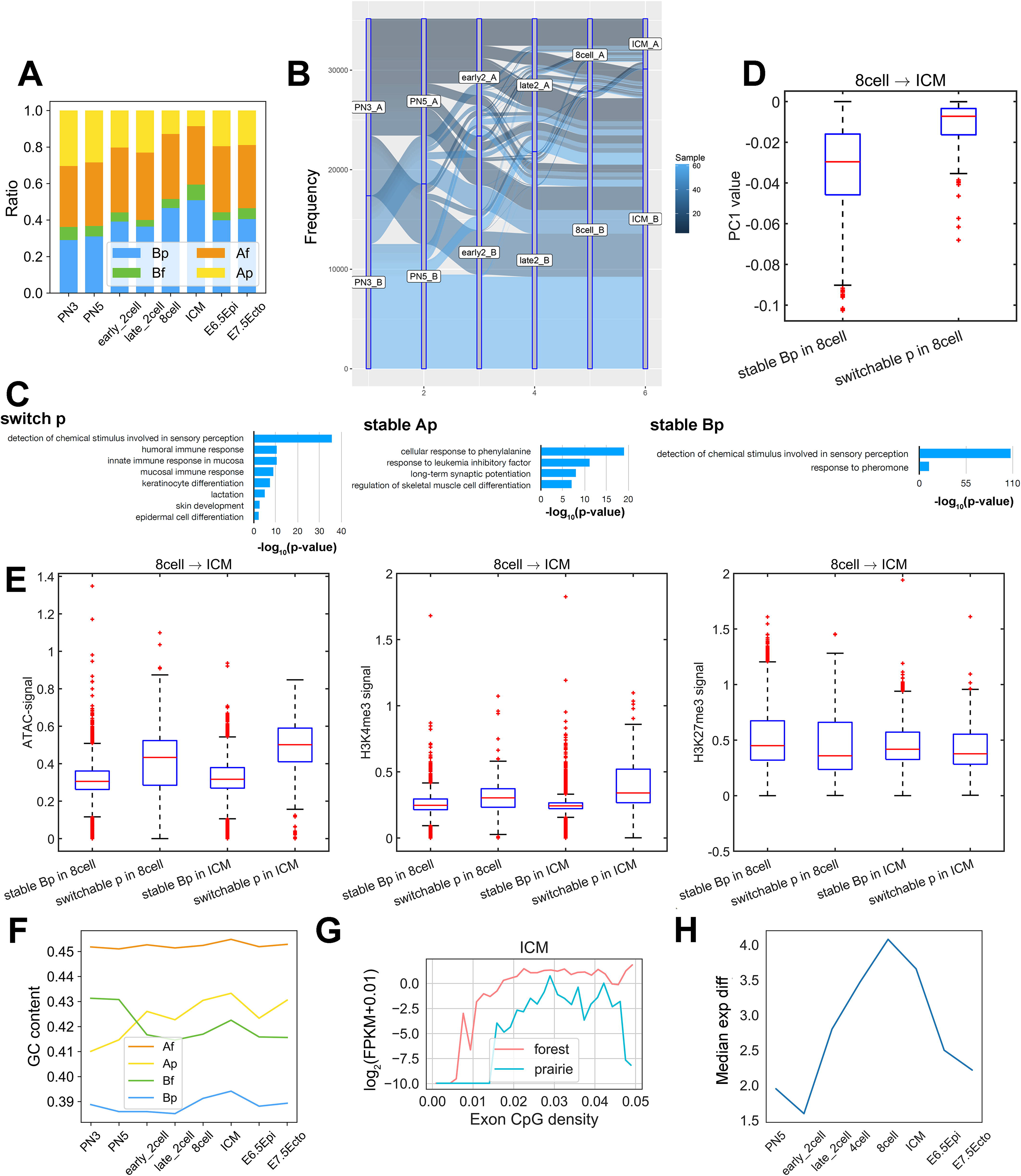
Compartment changes relate with domain segregation. **(A)** The proportions of four sequence components, Af, Bf, Ap, Bp (calculated according to the positioning of forests and prairies in compartments A and B), for each stage in the mouse embryonic development. **(B)** The dynamic partition of prairie into compartment A and B in mouse embryonic development. **(C)** Function annotation clustering of genes located in switch p, stable Ap and stable Bp regions, respectively. **(D)** The PC1 values of stable Bp (i.e., prairie regions constantly remaining in compartment B in both 8-cell and ICM) and switchable p (i.e., prairie regions located in compartment B in 8-cell while switching to compartment A in ICM) regions in 8-cell. **(E)** From 8-cell to ICM, the ATAC-seq, H3K4me3 and H3K27me3 signals of stable Bp and switchable p regions (which were explained in Figure 2D) in 8-cell and ICM. **(F)** The average GC contents of the four components in mouse embryonic development. **(G)** The relation between gene expression level and gene exon CpG density. Each data point represents the median expression level of one group of genes, the exon CpG density of which falls into one interval. **(H)** The expression difference between forest and prairie genes at different stages during mouse embryonic development.

Since the genome compartmentalization is weak at early stages of development(Du et al., 2017), for the robustness of compartment partition, we calculated the distribution of compartment index (C-index, a parameter to quantify the degree of compartmentalization, a larger value of which corresponds to a more compartment A-like environment, see “Methods”) at each stage for compartment A and B, respectively. The gradually increased discrepancy of C-index between compartment A and B supports the global structure segregation from zygote to ICM (Figure S3B). Then, we identified compartment A regions with positive C-index as strict compartment A (sA), B with negative C-index as strict compartment B (sB), and calculated the length of genome located in sAf (forest regions in sA), sAp, sBf and sBp, respectively. The result showed a trend similar to Figure 2A, that is, the proportion of prairies in strict compartment B (sBp) changes in the same trend as the chromatin segregation level, which rules out the possibility that the growth of Bp is an artifact due to compartment definition (Figure S3C).

In comparison with stable Bp regions, the prairie domains that switch the compartment type are particularly interesting as they may be regarded as mediators of nuclear architecture establishment during development. The analysis of compartment transformation during preimplantation development shows that 7.7% of prairies belong to compartment A (stable Ap), 26.3% of prairies always reside in compartment B (stable Bp), and the left 66.0% switches compartment type at least once (switchable p) (Figure 2B). Genes in the stable Ap and switchable p regions are enriched in immune- and ectoderm-related functions, while those in the stable Bp regions are not (Figure 2C). Previous analyses showed that forest and prairie domains tend to spatially segregate (and mainly correspond to compartment A and B, respectively), but to different extent in different cells(Liu et al., 2018). Moreover, forest-prairie spatial intermingling is cell-type specific, which is thought to be associated with prairie tissue-specific gene activation and the establishment of cell identity. Therefore, compartment B (heterochromatin) provides a silent environment for prairie genes, which wait to be activated through spatial interactions with forest/compartment A in the following differentiation stages (as seen in their spatial contact and expression properties in differentiated cells(Tian et al., 2020)). The function of genes in the stable Ap and switchable p regions thus supports that the gene expression in prairie domains plays an important role in cell fate determination. Indeed, our earlier analysis shows that prairie genes in compartment A are highly tissue-specific and by examining the functions of the related genes, one can deduce the tissue type of the associated sample(Liu et al., 2018). Besides, compared to forest genes, the expression of genes in prairie domains are more likely to be highly correlated with the compartment environment they reside in(Tian et al., 2020).

Next, to further investigate the differences between switchable prairie and stable prairie regions, we compared the chromatin features of these two kinds of regions between two adjacent stages. The cells we analyzed here include PN5, early 2-cell, late 2-cell, 8-cell and ICM due to the availability of Hi-C data. For each two adjacent stages (i.e., PN5 vs early 2-cell, early 2-cell vs late 2-cell, late 2-cell vs 8-cell, 8-cell vs ICM), we identified stable Bp regions and switchable p regions (p regions located in compartment B in the earlier whereas switched to compartment A in the later stage). Although these two kinds of prairie regions are both located in compartment B in the cell of the earlier stage, the PC1 value of switchable p regions was significantly more positive than stable Bp regions (Figures 2D and S3D) and accordingly, switchable p regions are significantly closer to A-B compartment boundaries (Figure S3E), indicating that Bp regions near the A-B boundary are more likely to switch. The analyses on ATAC-seq signal, H3K4me3 and H3K27me3 also support this observation (Figure 2E). For example, from 8-cell to ICM, the ATAC-seq and H3K4me3 signals of switchable p regions are significantly higher than stable Bp regions in both 8 cell and ICM, while the H3K27me3 signal of switchable p regions is weaker than stable Bp regions.

To further understand how DNA sequence affects chromatin compartmentalization in embryo development, we calculated the GC contents and CpG densities during development for the four genomic components Af, Bf, Ap and Bp (Figures 2F and S3F). Notably, the GC content of Ap changes in the same trend as the segregation degree in both mouse and human embryo development (Figures 2F and S3F). Since the proportion of Bp and the segregation level are highly correlated, when the degree of segregation increases, a higher portion of prairie domains partition into compartment B, leaving the remaining prairie sequences in compartment A more “bivalent” (low GC content regions of forests or high GC content regions of prairies(Liu et al., 2018)). Moreover, for mouse, both forest components (Af and Bf) have higher GC contents than prairie components (Ap and Bp) before the 2-cell stage, whereas compartment A DNA (Af and Ap) have higher GC contents than compartment B (Bf and Bp) after one passes the 2-cell stage (Figure 2F). The CpG densities vary in a similar trend (Figure S3F), which indicates that compartmentalization becomes more prominent after the 2-cell stage in mouse embryos, with higher-order structures being gradually established.

To investigate how domain segregation and CpG density affect gene expression, we calculated the median expression level of genes in forests and prairies as a function of CpG density. It can be seen from Figures 2G and S3G that in general the median gene expression level increases with the increase of CpG density in a certain range. Interestingly, for genes with similar exon CpG density, those in forests tend to show a higher expression than those in prairies (Figures 2G and S3G), showing the importance of large-scale DNA sequence property in gene expression (given that forest regions possess significantly higher CGI and thus CpG densities than prairies, the surrounding sequence environment of forest genes is thus CpG richer than prairie genes, suggesting a connection between their neighboring sequences and expression level of genes). Furthermore, the expression difference between forest and prairie genes (that is, the averaged difference between the two curves shown in Figure 2G for ICM and Figure S3G for other cells) increases from early 2-cell to 8-cell, then decreases from 8-cell to E7.5 ectoderm (Figure 2H), which is accordant with the chromatin segregation behaviors along the early embryo development.

### The Association between Domain Segregation and DNA Methylation

In the earlier study we showed that differences in the methylation levels between forests and prairies correlate well with chromatin spatial packing(Liu et al., 2018, 1). In the following, we analyzed methylation data for early embryonic cells obtained by four different research groups (see Additional file 2: Data sources). These data all show that the open sea (defined as the genomic regions excluding CGIs, CGI shores and CGI shelves(Sandoval et al., 2011)) methylation differences between forest and prairie domains in different cell types correlate well to their corresponding chromatin structural segregation behaviors (Figures 3A and S4A-S4C). During embryonic development, the absolute value of MDI (F-P open sea methylation difference index, see “Methods”, domain possessing a positive (negative) value indicates the methylation level of this domains is generally higher (lower) than its two flanking domains, therefore higher absolute value indicates the larger methylation level difference) increases from 2-cell to ICM stages, decreases at the ICM to E6.5 epiblast stage, and increases again in the further development to the E7.5 stage (especially for the E7.5 endoderm stage, Figure 3A). The variation of DNA methylation difference resembles that of the chromatin structural segregation. The correspondence between methylation difference and segregation degree during embryonic development further supports a connection between this epigenetic mark and the chromatin structure. In fact, such a correlation might have a simple explanation: Since forests contribute dominantly to the more accessible chromatin regions, they are presumably more prone to both DNA demethylation and methylation than prairies.

**Figure 3.**
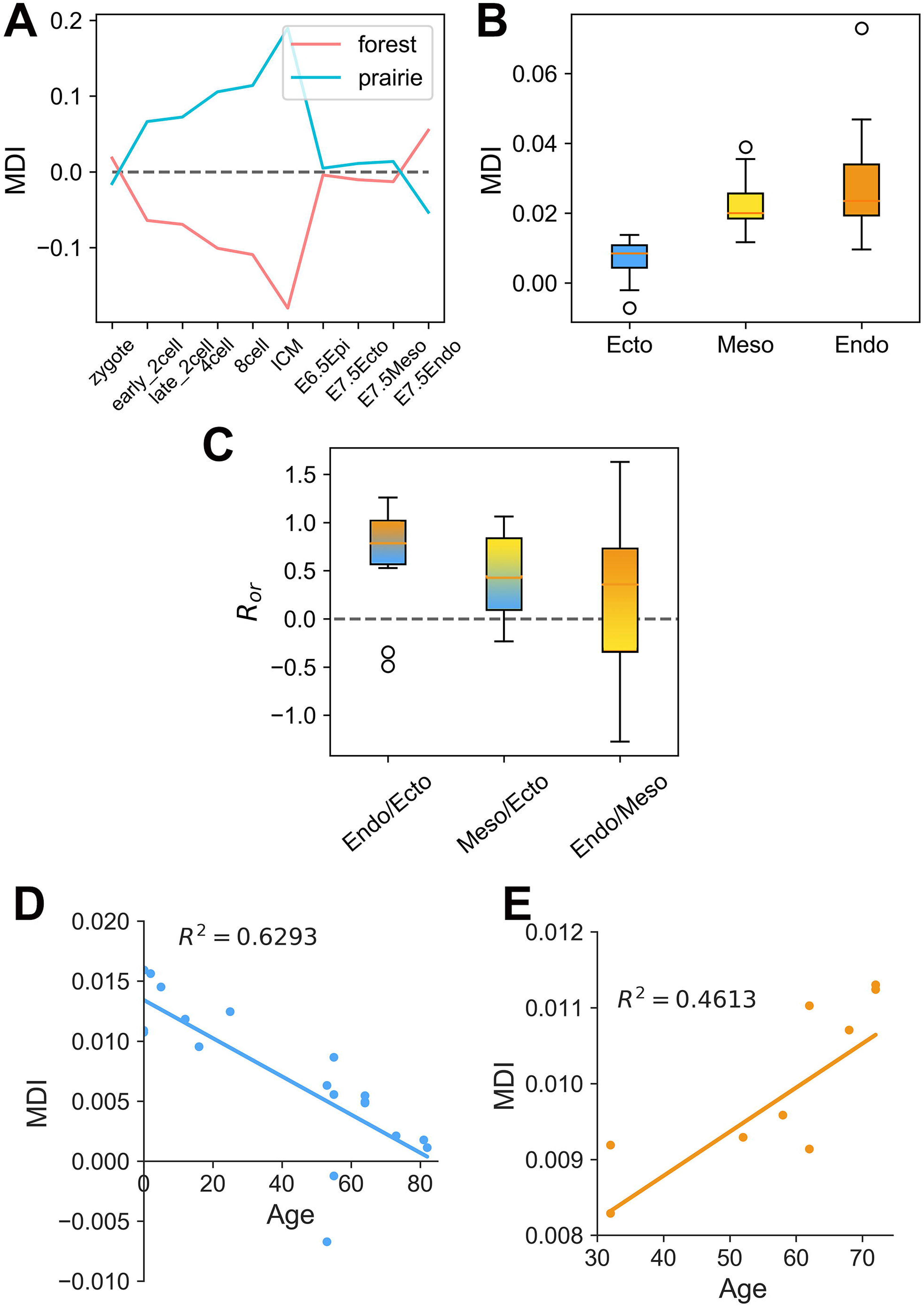
The association between domain segregation and DNA methylation. **(A)** The average forest/prairie open sea methylation level difference (MDI) at different stages during embryonic development. **(B)** The box plots of average MDI for human differentiated cells originating from endoderm, mesoderm and ectoderm, respectively, for forest domains. **(C)** The box plots of overall relative segregation ratio*R*_*or*_ (the definition can be seen below) of one tissue over another for forests. The three categories are tissues derived from endoderm and mesoderm over those derived from ectoderm, and endoderm over mesoderm. The parameter *R*_*or*_ was used to evaluate the differences of domain segregation behaviors between two tissues. For example, as for tissues 1 and 2, *N*_12_ and*N*_21_ represent the number of domains showing higher *DR*_*s*_ in tissues 1 and 2, respectively. Overall relative segregation ratio for tissues 1 and 2 is then defined as 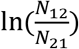, a positive value of which thus indicates the chromatin structure of tissue 1 is more segregated, compared to tissue 2. **(D, E)** The scatter plots of age and methylation difference for brain cells (D) and mature neutrophil cells (E) for forest domains.

Following such an observation, the extent of domain segregation of three germ layers can be inferred from the DNA methylation difference between the forest and prairie domains. As can be seen from Figure 3A, the calculated absolute value of MDI for endoderm is greater than that for mesoderm (The corresponding value of ectoderm is slightly less than mesoderm). This result suggests that the forest-prairie domains of endoderm is more segregated than those of mesoderm and ectoderm, which is indeed accordant with the analysis on Hi-C data (unpublished results). Interestingly, domain segregation levels of the different tissues can be seen to reflect the germ layers they originate from, as shown below from the analyses of both DNA methylation and structural data.

It was found based on the Hi-C data that ectoderm-derived cortex chromatin is less segregated than that of endoderm-derived liver(Liu et al., 2018). Since the above analysis suggests chromatin in ectoderm to be less segregated than endoderm (Figure 3A), we decided to examine whether chromatin structures of differentiated tissues also show such a trend, therefore exhibiting a germ layer dependence. We analyzed average methylation difference levels of 138 mouse differentiated cells and 45 human differentiated cells (38, 34, 66 cells originate from endoderm, mesoderm and ectoderm respectively for mouse and 13, 19, 13 for human, see Additional file 2: Data sources)(The ENCODE Project Consortium, 2012; Lister et al., 2013; Schultz et al., 2015). As shown in Figure 3B, the absolute values of MDI for forest domains for human ectoderm-derived tissues are significantly smaller than those for mesoderm-(*p* = 4.938 × 10^−7^ by two-sample t-test) and endoderm-derived (*p* = 5.696 × 10^−4^ by two-sample t-test) tissues, in the same as the segregation degree of the three embryonic germ layers. The data on prairie domains of human tissues show the same trend, so do those on both forest and prairie domains of mouse tissues (Figure S4D-S4F). To further validate this finding, we analyzed the Hi-C data of 14 human tissues(Schmitt et al., 2016) (see Additional file 2: Data sources) and compared the segregation ratio of these tissues derived from different germ layers. To quantify the difference between two tissues, we defined and calculated the overall relative segregated ratios (*R*_*or*_) of one tissue over another, a positive value of which represents that chromatin in the former is more segregated than that in the latter (see “Methods”). For tissues derived from endoderm and mesoderm, their overall relative segregated ratios over those originated from ectoderm (cortex and hippocampus) are generally positive (Figure 3C), indicating that chromatin of the former indeed tends to be more segregated than that of the latter. Similarly, the overall relative segregation ratios of tissues derived from endoderm over those derived from mesoderm also tend to be positive (Figures 3C and S4G). Together, these results are all consistent with the notion that the segregation level of a differentiated cell shows the corresponding germ layer signature, that is, the order of segregation degrees for the three germ layers is in accordance with that of differentiated tissues.

More interestingly, we found that the absolute values of MDI for brain decrease with aging, and the trend is opposite for mature neutrophil cells, as shown in Figures 3D and 3E. Earlier studies have found that DNA hypomethylation, which is more likely to occur in prairie domains, correlates with the cell cycles. Partially methylated domains (PMDs) in tumor cells are mainly composed of prairies(Xue et al., 2020) and PMD hypomethylation increases with age, which appears to track the accumulation of cell divisions(Zhou et al., 2018). Xuan Ming et al. also found that solo-WCGW sites display aging- and cancer-associated hypomethylation, which exhibit low maintenance efficiency during cell cycle(Ming et al., 2020). For chromatin structural changes, we compared the segregation ratio in G1 stage to that in the late S ~ G2 stage, and found that forest and prairie tend to become more separated in the late S ~ G2 stage (Xue et al., 2020). The enhanced segregation supports that mitosis is conducive to a more segregated chromatin structure. Therefore, we speculate that the different methylation patterns associated with aging among different tissues may reflect their different cell division patterns: the ectoderm-originated brain cells hardly divide, while liver cells constantly undergo cell cycles. The observed MDI differences between these cells are consistent with their different dividing patterns in life. Such a consistency makes an understanding of the mechanistic connection between methylation and cell-division patterns highly desired.

### ZGA and Associated 3D Genome Architecture Change

To further investigate how transcription is associated with chromatin structure, we analyzed the chromatin structure changes from normal early 2-cell to late 2-cell at different genomic distance scales, since ZGA typically occurs at this period. At small genomic distances (<500 kb), the F-P ratio for late 2-cell is larger than that for early 2-cell, while P-P ratio is smaller for the former (Figure 4A, upper two figures). Our previous work has revealed that prairie-forest intermingling was associated with prairie gene activation(Liu et al., 2018), therefore, we speculate that the increase of F-P ratio may be associated with ZGA. At large distances (500 kb-~20 Mb), the P-P ratio (F-P ratio) of late 2-cell was larger (smaller) than early 2-cell (Figure 4A, upper two figures), indicating that at such scale, the forest-prairie separation is enhanced, accordant with the increase of compartmentalization degree(Du et al., 2017) (given that compartment A and B are mainly composed of forest and prairie regions, respectively). Further, compared to early 2-cell, chromatin of late 2-cell establishes more spatial interactions over longer distances (>50 Mb, Figure 4A, down two figures). However, the compartmentalization is weak at this scale (compared to the F-P/P-P ratio of ICM in the same range (i.e., >50 Mb) and late 2-cell itself in small range (i.e., 50 Mb), Figure 4A, down two figures).

**Figure 4.**
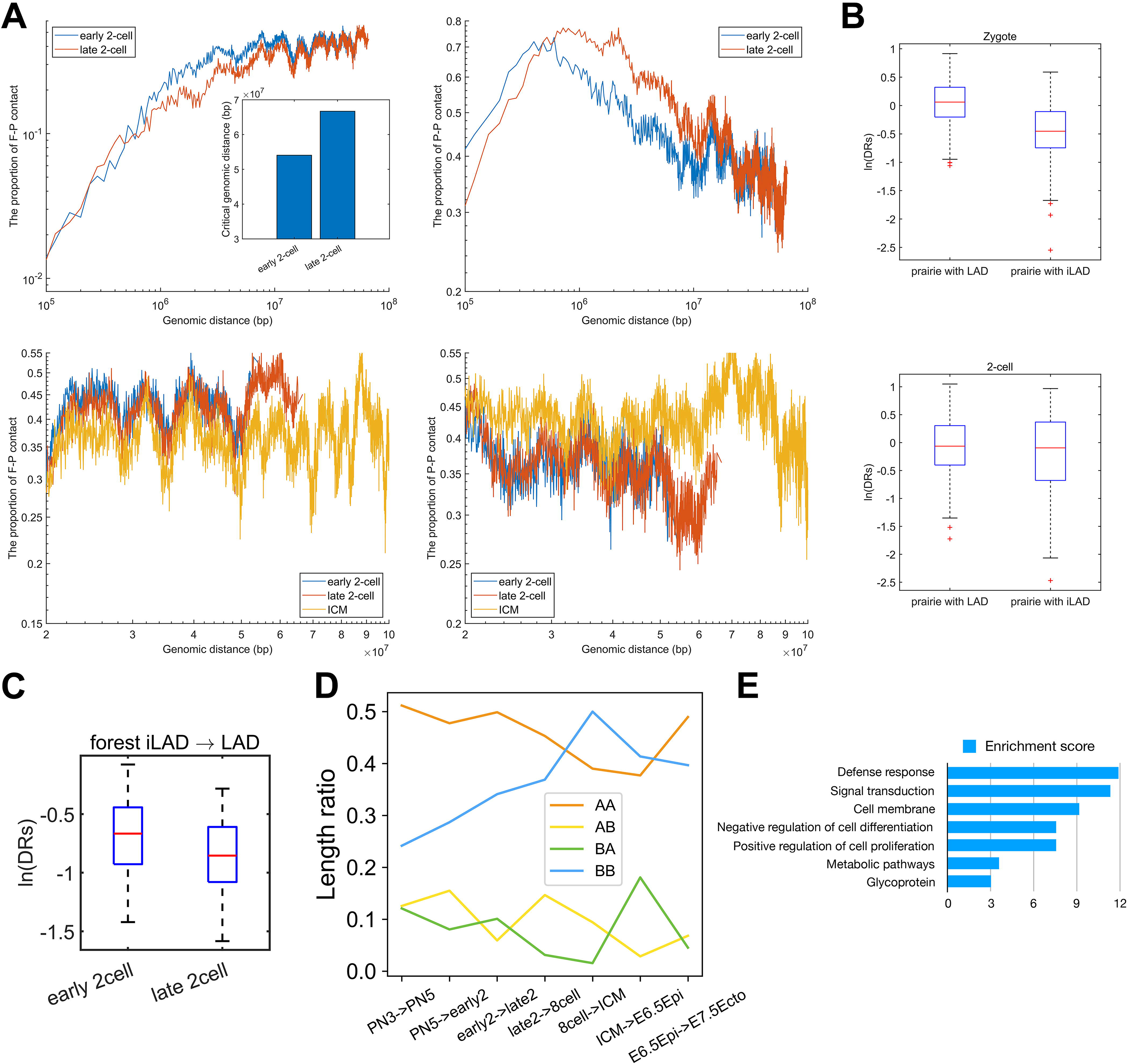
ZGA and associated 3D genome architecture change. **(A)** The upper two figures represent the proportion of F-P and P-P spatial interactions within the top 10% contact probabilities under one certain genomic distance, for normal early 2-cell, late 2-cell. The lower two figures were the amplifications of the partial regions of two upper figures and ICM was also included for comparison. Inner plot represents the critical genomic distance of early 2-cell and late 2-cell. **(B)** The domain segregation ratio, *DR*_*s*_, of prairie domains that are/are not in LAD regions. **(C)** The domain segregation ratio of forest domains switching from iLAD to LAD during ZGA, in early 2-cell and late 2-cell. **(D)** The proportion of four components (A→A, A→B, B→A, B→B) in each transition (e.g., from early 2-cell to late 2-cell). The length ratio of e.g., AA for PN3->PN5 indicates the ratio between the genome length of stable A regions (between PN3 and PN5) and the entire genome length. **(E)** Function annotation clustering of prairie genes switching from B to A during ZGA.

LADs, which resemble compartment B, are established quickly after fertilization. More than 75% of LAD domains are prairies across preimplantation development in mouse and the domain segregation ratio ( *DR*_*s*_) of prairie domains overlapping with LAD is significantly larger than those that are not (Figure 4B, *p* = 2.9 × 10^−34^ and 0.01 for zygote and 2-cell, respectively, Welch’s unequal variance test). We further analyzed the correlation between LAD mixing and prairie/forest mixing at the ZGA stage in mouse. LAD/iLAD (inter-LAD) regions in zygote and 2-cell stage were obtained from previous studies(Borsos et al., 2019). We then identified forest/prairie domains that become significantly more intermingled or segregated (i.e., *F*_*mix*_/*F*_*seg*_/*P*_*mix*_/*P*_*seg*_, see “Methods”) during ZGA. The number of forest domains switching from iLAD to LAD state is 139, half of which (70 domains) belong to *F*_*mix*_. In contrast, the proportion of *F*_*mix*_ domains within forest domains remaining in the iLAD state is only 24.8% (*p* = 1 × 10^−7^ by Fisher’s exact test). Accordingly, forests that change from iLAD to LAD regions show a significant decrease in domain segregation ratios (Figure 4C, *p* = 6.7 × 10^−5^ by two-sample t-test). Such a result indicates that the forests changing from iLAD to LAD gain contacts with prairies, and a correlation does exist between LAD mixing and forest mixing during ZGA.

We next examine how compartment switch is associated with ZGA. Firstly, we examined the compartment change during embryonic development. The overall length of genomic domains changing from compartment A to B are shorter than those switching from compartment B to A in ZGA (Figure 4D). Such a phenomenon was also observed in implantation (from ICM to E6.5Epi). In addition, the prairie genes that move from compartment B to A in ZGA are found to be rich in functional annotation of “defense response”, “signal transduction”, “cell membrane”, “negative regulation of cell differentiation” and “positive regulation of cell proliferation,” identified using DAVID (Huang et al., 2008, 2009) (see “Methods”, Figure 4E). These results indicate that the domain mixing during ZGA is heavily related to functions of cell proliferation.

### Implantation-related Domain Mixing in Differentiation

Since it was shown that the forest-prairie inter-domain interaction coincides with tissue-specific gene activation in differentiation(Liu et al., 2018), one wonders how the achievements of tissue-specific functions are reflected by domain mixing in implantation. We again calculated domain segregation ratio *DR*_*s*_ for each forest/prairie domain at related embryonic developmental stages (i.e., ICM and E6.5Epi), and observed that *DR*_*s*_ decreases for 89.0% of prairie domains at implantation (Figure 5A). To be more specific, we identified DNA domains that undergo significant changes of segregation, and denoted more strongly segregated and more mixed domains in forest and prairie as *F*_*seg*_, *P*_*seg*_, *F*_*mix*_, and *P*_*mix*_, respectively (see “Methods”). In the implantation stage, 71.0% of prairie domains belong to *P*_*mix*_, while 21.3% of forest domains become more mixed (Figure 5B).

**Figure 5.**
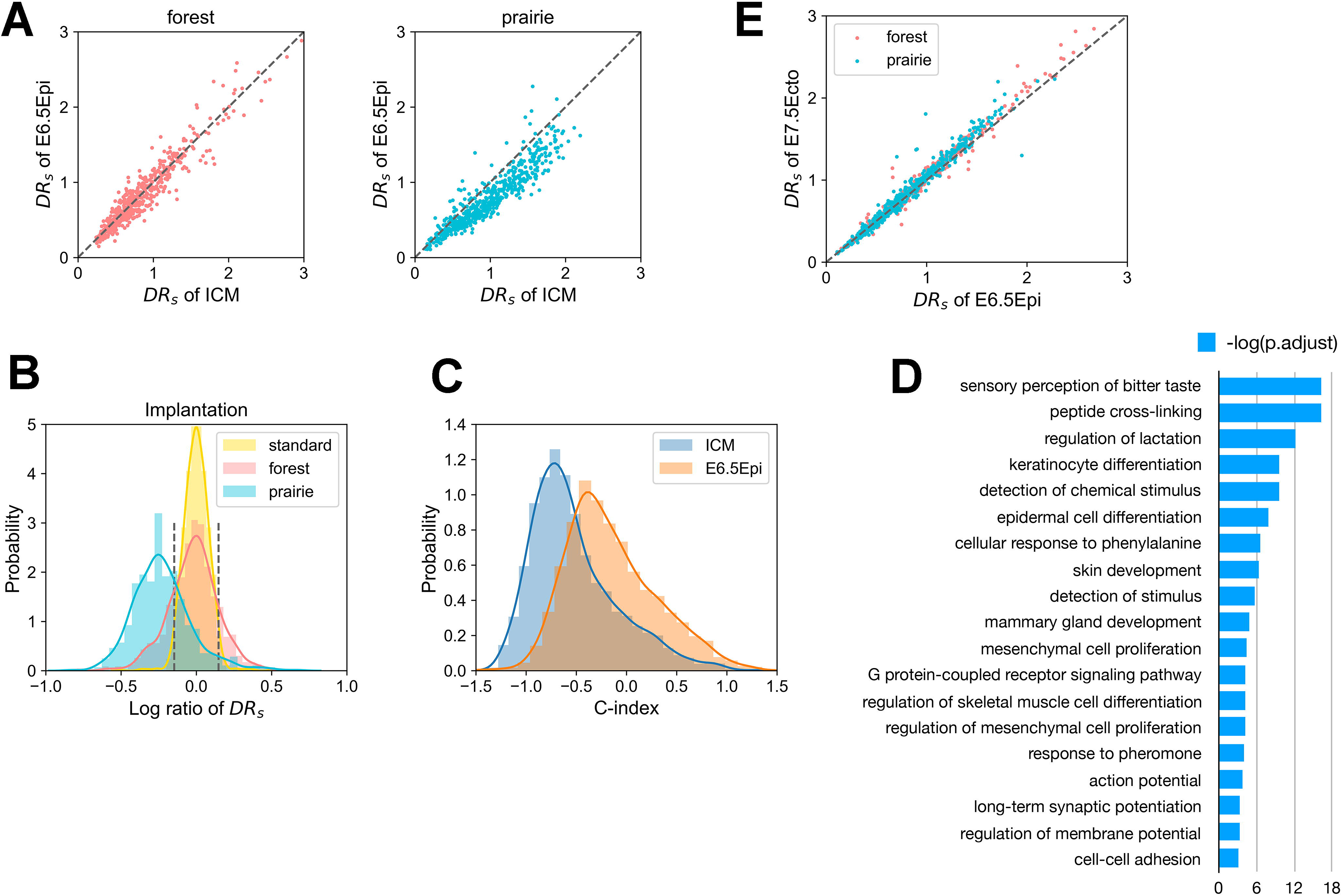
Implantation-related domain mixing in differentiation. **(A)** The scatter diagrams of domain segregation ratio *DR*_*s*_ for forests and prairies during implantation. **(B)** The distribution of the logarithm ratio of *DR*_*s*_ between E6.5Epi and ICM for forests and prairies. The plot, the legend of which is “standard”, represents the logarithm ratio of *DR*_*s*_ between two ICM replicates from the same lab. Significantly more strongly segregated domains are taken as those whose logarithm ratio of *DR*_*s*_ values exceed the 97.5^th^ percentile of the “standard” distribution, and accordingly, domains whose logarithm ratio of *DR*_*s*_ is smaller than the 2.5^th^ percentile of the “standard” distribution are regarded as significantly more mixed domains. The corresponding threshold values are 0.1469 and −0.1469, respectively, which was labeled using two black dotted lines. **(C)** The change of compartment index (C-index) for less-segregated prairies, *P*_*mix*_, from ICM to E6.5 epiblast stage. **(D)** Functional assignment of genes in the *P*_*mix*_ during implantation. **(E)** The scatter plot of *DR*_*s*_ for forests and prairies between E6.5 epiblast and E7.5 ectoderm.

As for *P*_*mix*_ regions following implantation, we first examined their structural changes by calculating the compartment index (C-index) changes, a larger value of which corresponds to a more compartment A-like environment (see “Methods”). Along with the ICM to E6.5 epiblast transition, 86.9% of *P*_*mix*_ domains have an increase of C-index (entering a more compartment A-like environment), implying that genes in these regions move to a more active environment (Figure 5C). We analyzed next the functional enrichment for genes located in the *P*_*mix*_ domains using ClusterProfiler(Yu et al., 2012) and DAVID(Huang et al., 2008, 2009) (see “Methods”). The associated *P*_*mix*_ genes are characterized by “sensory perception of taste”, “regulation of lactation”, “keratinocyte differentiation”, “epidermal cell differentiation”, “epidermis development”, “skin development”, “mammary gland development” and “long-term synaptic potentiation” (Figure 5D). Interestingly, these terms are related to mammary glands, epidermis and nervous system, all of which are differentiated from ectoderm(Li et al., 2013), indicating that the mixing of prairie domains with forests during implantation is highly pertinent to ectoderm differentiation. A supporting observation is that chromatin structures of E6.5 epiblast and E7.5 ectoderm are similar as seen by their similar *DR*_*s*_ values (Figure 5E). Such an observation suggests that chromatin enters an ectoderm-like state shortly after implantation, consistent with earlier studies which showed the primed ESCs, representing a post-implantation epiblast cell state, to subsequently develop towards the ectoderm(Ji et al., 2016). In addition, during the implantation process, the expression level of 309 prairie genes increases, while 162 prairie genes show decreased expression level. Here we further found that prairie domains harboring genes with elevated expression levels tend to become more forest-prairie mixed from ICM to E6.5Epi, compared to genes with lowered expression (Figure S5A, *p* = 4.163 × 10^−4^ by two-sample t-test), consistent with our previous finding on the vital role of forest-prairie spatial interactions in prairie gene activation(Liu et al., 2018). Moreover, we found that *P*_*mix*_ genes with increased expression tend to be enriched in neuro-related functions (Figure S5B), indicating that neuroectoderm may differentiate ahead of epidermal ectoderm. Finally, to confirm the roles of domain segregation change on biological functions, we analyzed the structural changes of domains containing lineage-specific genes. We identified 552 lineage-specific genes for E6.5 epiblast stage (see “Methods”, i.e., genes specifically and highly expressed in E6.5 epiblast stage), among which 474 genes locate in forest domains and 78 genes in prairie domains. Ninety-seven percent of the prairie lineage-specific genes located in *P*_*mix*_, suggesting that the mixing of prairies into forests does associate with lineage specification during implantation. Meanwhile, the prairie genes that move from compartment B to A during implantation are also functionally enriched in ectoderm differentiation and utero embryonic development (Figure S5C), in accordance with above analysis on *P*_*mix*_ regions.

## DISCUSSION

### Sequence-based Chromatin Domain Segregation

In the present study, we described the chromatin structural changes during embryonic development from zygote to the post-implantation stages and, in particular, how DNA sequence affects the chromatin structural changes. We also investigated the relationship between DNA domain segregation and the genetic/epigenetic properties. Earlier studies have shown gradual compartmentalization in embryonic development(Du et al., 2017). Here we connect the 3D structure change to DNA sequence properties and perform functional analysis of these structural changes. The sequence-structure-function analysis provides a common-ground for the understanding of different biological processes. In particular, two types of genomic domains defined earlier, forests and prairies, are shown to generally undergo an enhanced spatial separation in early embryo development of both mouse and human, which is also reflected in the DNA methylation and gene expression difference between the two domain types. Interestingly, noticeable domain mixing does occur at two important stages, ZGA, when zygotic genes begin to express, and implantation, during which differentiation starts to form germ layers. During ZGA process, short-range (<500 kb) forest-prairie spatial interactions increase, which was thought to be associated with gene activation since our previous work has revealed that prairie genes tend to move to a more forest environment for activation (Tian et al., 2020). Besides, we also investigated in detail the chromatin structural changes during ZGA process at different scales. During implantation, we observed a conspicuous decrease of segregation ratio *R*_*s*_(*d*) for prairies at large genomic distances (>3 Mb). Intriguingly, almost all E6.5 epiblast-specific prairie genes resided in *P*_*mix*_ domains, again emphasizing the intimate link between domain mixing and gene activation (lineage specification). Genes in the prairie domains that become more mixed in implantation are prominently related with ectoderm differentiation.

Analysis of Hi-C data in differentiated and senescent cells (Liu et al., 2018), showed a consistently enhanced segregation of chromatin structures, with a gradual establishment of long-range DNA contacts from zygote to senescence. At the zygotic stage, few high order structural features exist and the chromatin is organized similarly to a random coil (Figure 6A). As the embryo develops, local structures (loops and TADs) become more prominent and the two different types of domains segregate from each other to form compartments. Such a trend continues through ICM (Figure 6B). After implantation and as differentiation starts, a subset of prairie domains tends to mix into the active environment, activating associated genes (Figure 6C). During senescence, prairie domains congregate further, some of which detach from nuclear membranes and cluster inside the nucleus, permitting long-range contacts establishment(Chandra et al., 2015) (Figure 6D). Previous analysis showed that chromatin domain segregation continues in cell differentiation and senescence(Liu et al., 2018, 1). It appears that the overall trend of chromatin structure change follows an establishment of long-range contacts from birth to senescence, as seen from the segregation level change (Figures S6A-S6C). In this process, intra-domain contacts are established first, followed by long-range inter-domain contact formation, and then the establishment of lineage-specific inter-domain contacts, which relates with cell differentiation (Figures S6D-S6G).

**Figure 6.**
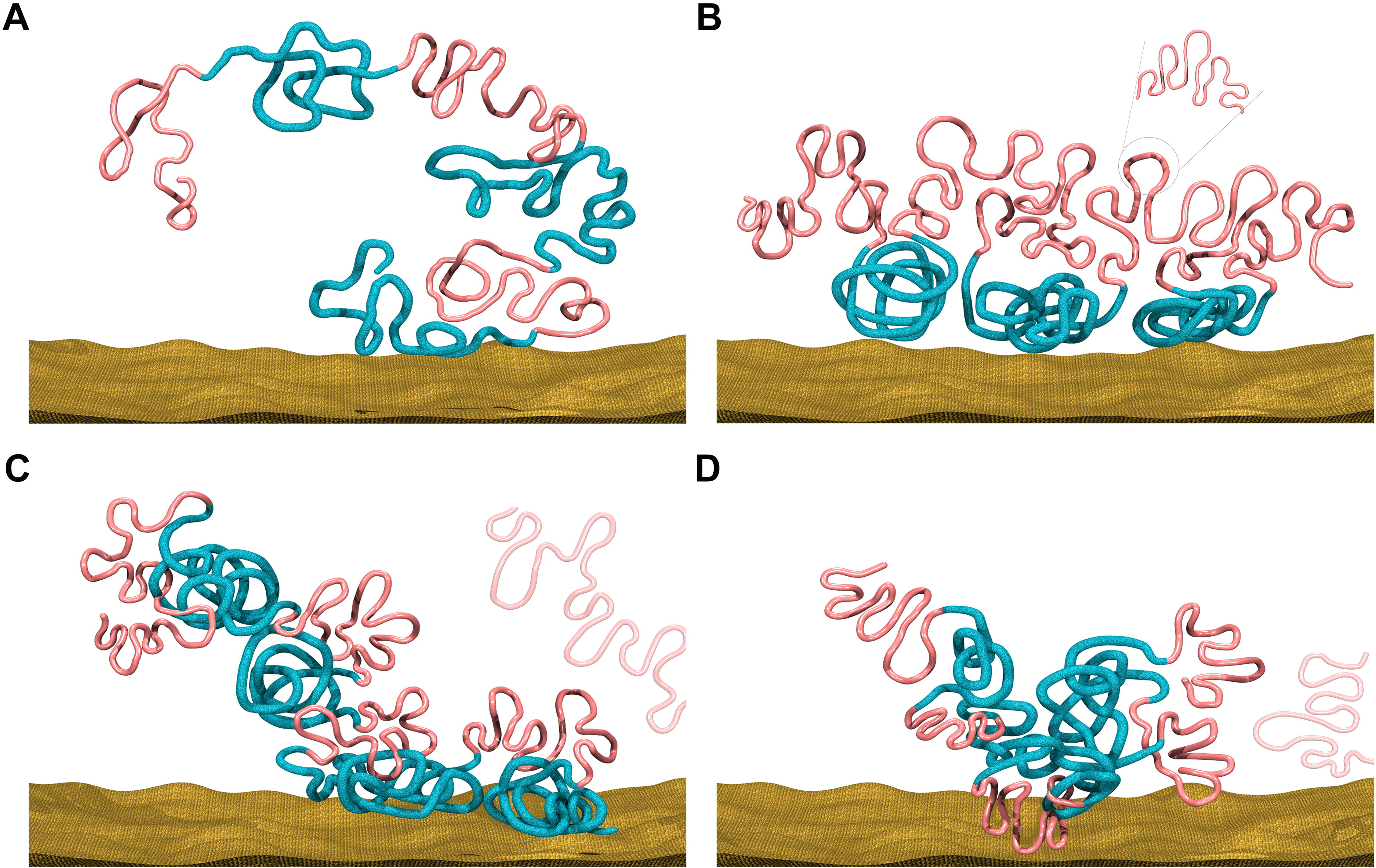
Schematic pictures of chromatin structural patterns at different stages. **(A)** In the beginning zygotic stage, chromatin has little long-range structural features. Here the red lines represent CGI forest domains, and the blue ones represent CGI prairie domains. **(B)** In ICM stage, the two different types of domains segregate from each other to form compartments. **(C)** In differentiated cells, prairies containing tissue-specific genes tend to form contacts with forests. **(D)** During senescence, prairies congregate further (with an increased probability to detach from the nuclear membrane).

### The Association between Transcription Inhibition and Genome Architecture

To investigate how transcription inhibition affects chromatin architecture, we compared the F-P and P-P ratio within the top 10% contact probabilities under one certain genomic distance between PN3, normal late 2-cell and *α*-amanitin-treated-20h cell (*α*-amanitin-treated late 2-cell). The critical genomic distance of *α*-amanitin-treated-20h cell is significantly smaller than that of normal late 2-cell, and the F-P and P-P ratio of *α*-amanitin-treated-20h cell are between PN3 and normal late 2-cell (Figure S7A), indicating that transcription inhibition can slow down the establishment of long-range contact and forest-prairie separation. We further compared the F-P and P-P ratio between *α*-amanitin-treated-20h cell and *α*-amanitin-treated-45h cell, and found that the critical genomic distance and P-P ratio of the latter are larger than the former, the F-P ratio of the latter is smaller than the former (Figure S7B), hinting that when transcription is inhibited, long-range chromatin contacts can still be established and forest-prairie separation also progresses, although at a reduced rate.

### Other Factors that May Relate to Chromatin Structure Change

Sequence-based chromatin domain segregation suggests that factors effecting domain interactions could lead to functional chromatin structure changes. Besides signaling and transcriptional factors, physical parameters could also be important. Interestingly, a correlation seems to exist between chromatin domain segregation and body temperature during early embryonic development. For human, the variation of BBT (basal body temperature) corresponds closely to the change in domain segregation. Firstly, BBT rises 0.5~1°F immediately after ovulation and remains high throughout the luteal phase, even after implantation(Su et al., 2017). The segregation ratio *R*_*s*_ increases along with the development from zygote to ICM, except for the early to late 2-cell stage, which coincides with a small temperature dip nearly two days after ovulation, nicely corresponding to the mixing of chromatin domains observed at this stage. Moreover, a one-day implantation dip of BBT occurs nearly one week after ovulation. Correspondingly, a significant domain mixing is observed at this stage of development, from ICM to E6.5 epiblast. As widely known, implantation dip is effective in detecting pregnancy in humans(Refinetti and Menaker, 1992) and continuous BBT measurement reliably allows for the earliest and most accurate detection of pregnancy(Smarr et al., 2016). For mouse, researchers did find that an ultimately-successful pregnancy is characterized by significantly more min/day of relative low CBT (core body temperature) in the second day after mouse pairing which rightly corresponds to ZGA, than for pregnancies which subsequently terminated prematurely(Smarr et al., 2016). These observations give rise to the possibility that temperature shifts are closely coupled with large scale chromatin structural changes during embryonic development, in accordance with high temperatures to be favorable for segregation of the prairies, and low temperatures, for a more forest-prairie mixed chromatin structure(Liu et al., 2018; Quan et al., 2020).

Besides human and mouse, temperature is also found to be important in embryonic development for many other species. One interesting example is the BBT of hibernating bears. The body temperature of the pregnant female bears decreases at the implantation stage, but remains 3-4 degrees higher and significantly more stable than non-pregnant females or males(Friebe et al., 2014), indicating the importance of a high and steady temperature in bear embryo development. In addition, researchers also reported chromosome de-condensation in rice seedlings underwent cold stress(Liu et al., 2017). All these results are consistent with the postulation that an increase of temperature correlates with an enhanced chromatin domain segregation, and imply that temperature could be a non-negligible factor influencing global chromatin structure, thus a possible parameter for affecting chromatin reprogramming or modification.

The domain segregation of the earliest fate-committed cell types also shows correlations with those of the differentiated cells. Tissues differentiated from endoderm generally have more strongly segregated chromatin structures than those from ectoderm. Such a difference between ectoderm and endoderm can be seen as early as from an E7.5 embryo. As another example, the tissue temperatures of interscapular brown adipose tissue (IBAT), liver, and the cortical (T_IBAT_, T_L_, T_CO_, respectively) were found to follow such an order: T_CO_ ≪T_IBAT_ < T_L_(Dewasmes et al., 2003). This order of temperature values is consistent with that of the corresponding chromatin domain segregation level, which increases in the order of cortical, IBAT and liver. Furthermore, these tissues are known to originate from ectoderm, mesoderm and endoderm, respectively, also consistent with the degree of domain segregation of the different germ layers. Although much more data is needed and many more factors such as protein structures and protein-DNA interactions are likely involved in affecting the chromatin structure(Hnisz et al., 2017), the cause of such correlations is worth of more detailed investigations.

Cell cycle is another important factor that could influence chromatin domain segregation. During embryonic development, the establishment of TAD structure was reported to require DNA replication but not zygotic transcription(Du et al., 2017; Ke et al., 2017), implying the critical role of cell division in 3D structural formation. In fact, our analysis shows that different cell division behaviors of different tissues including those derived from different germ layers further increase their discrepancy in structural segregation as well as in methylation pattern. The current and earlier analyses(Liu et al., 2018) indicate that the DNA methylation are in affect associated closely to chromatin conformation. The age related hypomethylation appears to track closely the accumulation of cell divisions(Zhou et al., 2018). We speculate that along with hypomethylation (leading to the formation of partially methylated domains), prairie domains tend to segregate more stably in the less active spatial domains of heterochromatin(DEEP Consortium et al., 2018).

## CONCLUSION

In this study, we delineated chromatin structural and epigenetic reprogramming in early embryonic development based on the sequence-based domain segregation model, and correlated the overall chromatin structural change with functional implementation in two vital processes, ZGA and implantation. Genes within less segregated prairie domains during implantation show lineage-relevant functions and thus may play important roles in corresponding processes. Therefore, detailed analyses on gene expression and regulation of transcription factors are in need for understanding the related mechanisms.

## DATA AVAILABILITY STATEMENT

All data analyzed during this study are publicly available. The detailed data information can be found in Additional file 2 Data Sources.

## ACKNOWLEDGEMENTS

We would like to thank Prof. Xiaoliang S. Xie for helpful discussions, Zhenhai Du for the initial processing of Hi-C data, and Yupeng Huang for helping us to draw figure 6.

## AUTHOR CONTRIBUTIONS

Conceptualization, Y.Q.G., H.Q., H.T. and S.L.; Methodology, Y.Q.G., H.Q., S.L. and H.T.; Software, H.Q., S.L., Y.X. and H.T.; Validation, W.X. and Y.Z.; Formal Analysis, H.Q., S.L. Y.X. and H.T.; Investigation, W.X. and Y.Z.; Writing-Original Draft, H.Q. and H.T., Writing-Review & Editing, Y.Q.G., S.L., H.T., Y.Z. and W.X.; Visualization, H.Q., X.Y. and H.T; Supervision, Y.Q.G.; Funding Acquisition, Y.Q.G.

## SUPPLEMENTARY MATERIALS

### Additional file 1

Figures S1-S7.

### Additional file 2

Tables S1 provides a summary of all used data.

Table S2-8 contains information of analyzed mouse Hi-C data, human Hi-C data, mouse embryo methylation data, mouse tissue methylation data, human tissue methylation data, mouse expression data and mouse epigenetic data, respectively.

### Additional file 3

CGI forest definition: 1cluster_by_nearby_dist_m.m;
distance-dependent segregation ratio calculation: 2Rs(d)_calculation.m;
MDI calculation: 3mm9_embyo_diff_pf_os.m;
overall and stringent relative segregation ratio:4overall_stringent_Rs.m;
domain segregation ratio calculation: 5DRs_calculation.m;
compartment index calculation: 6Comp_index_implan.m

