## Supplemental file for "Progressive domain segregation in early embryonic development and underlying correlation to genetic and epigenetic changes"

### Supplementary Figures

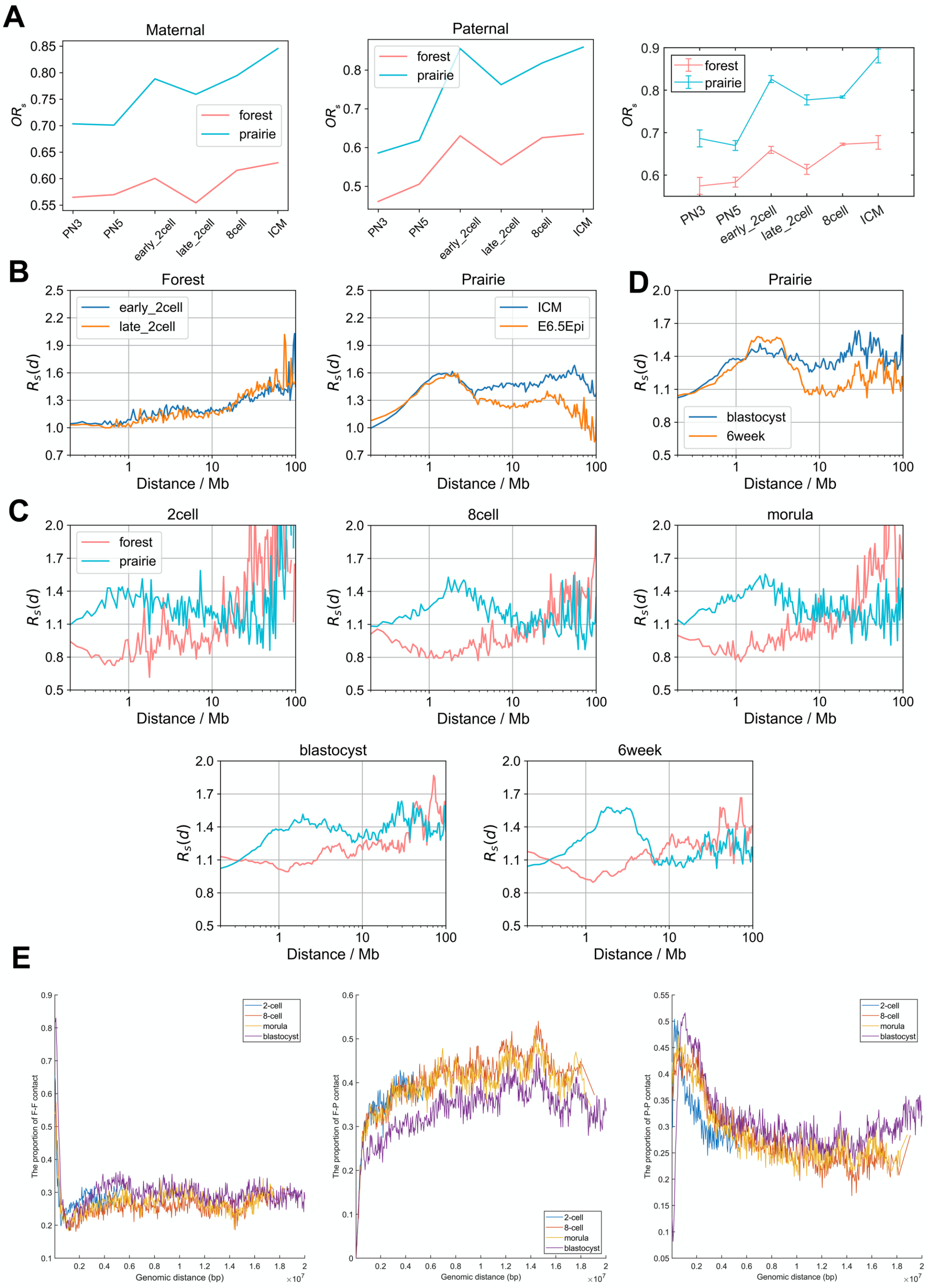


**Figure S1. Domain segregation in early embryonic development.** Related to Figure 1.

(A) The overall segregation ratio of forests and prairies at each stage in mouse embryonic development for maternal (left figure) and paternal chromatins (middle figure). The $OR_s$ generally increases when embryonic cells develop from zygote to ICM, while one dip occurs at the early to late 2-cell stage, both for maternal and paternal chromatins. Besides, data from different samples (right figure, for instance, there are 12 samples in PN5, such as PN5_rep1, PN5_rep4, PN5_maternal_rep1, PN5_paternal_rep1 and so on) are also analyzed. Error-bar: Mean±SE (standard error).

(B) The comparisons of distance-dependent segregation ratio $R_s(d)$ between different stages in mouse. The $R_s(d)$ of forests at 1~10 Mb for early 2-cell is slightly smaller than that for late 2-cell. At distances larger than 3 Mb, the $R_s(d)$ of prairies for ICM is much larger than that for E6.5 epiblast in mouse.

(C) The distance-dependent segregation ratio $R_s(d)$ of forest and prairie domains at each stage in human embryonic development.

(D) The comparisons of distance-dependent segregation ratio $R_s(d)$ between blastocyst and 6week in human. The $R_s(d)$ of prairies for blastocyst is significantly larger than that for the 6-week stage in human.

(E) The proportions of F-F, F-P and P-P spatial interactions within top 10% contact probabilities under one certain genomic distance at each stage in human embryo development.

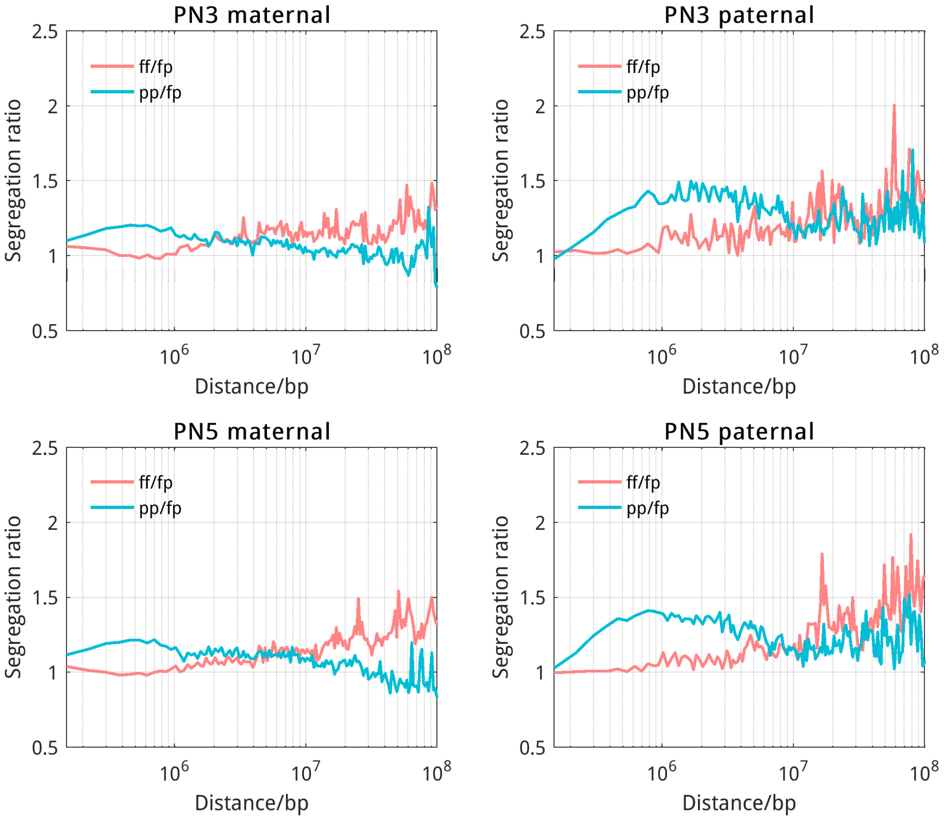


**Figure S2. Comparisons between maternal and paternal genomes.** As the calculation of $OR_s$ uses contacts without distance normalization, $OR_s$ is mainly influenced by interactions at short genomic distances. A better indicator would be contact ratio at the length scale of compartments (~Mb). To compare segregation degrees at all genomic lengths, we calculated segregation ratios along genomic distances, and found that the maternal genome is less segregated than the paternal genome at the ~300 kb to several Mb region. The difference between the maternal and paternal chromatin segregation is thus in accordance with lack of compartments in maternal genome compared to paternal one.

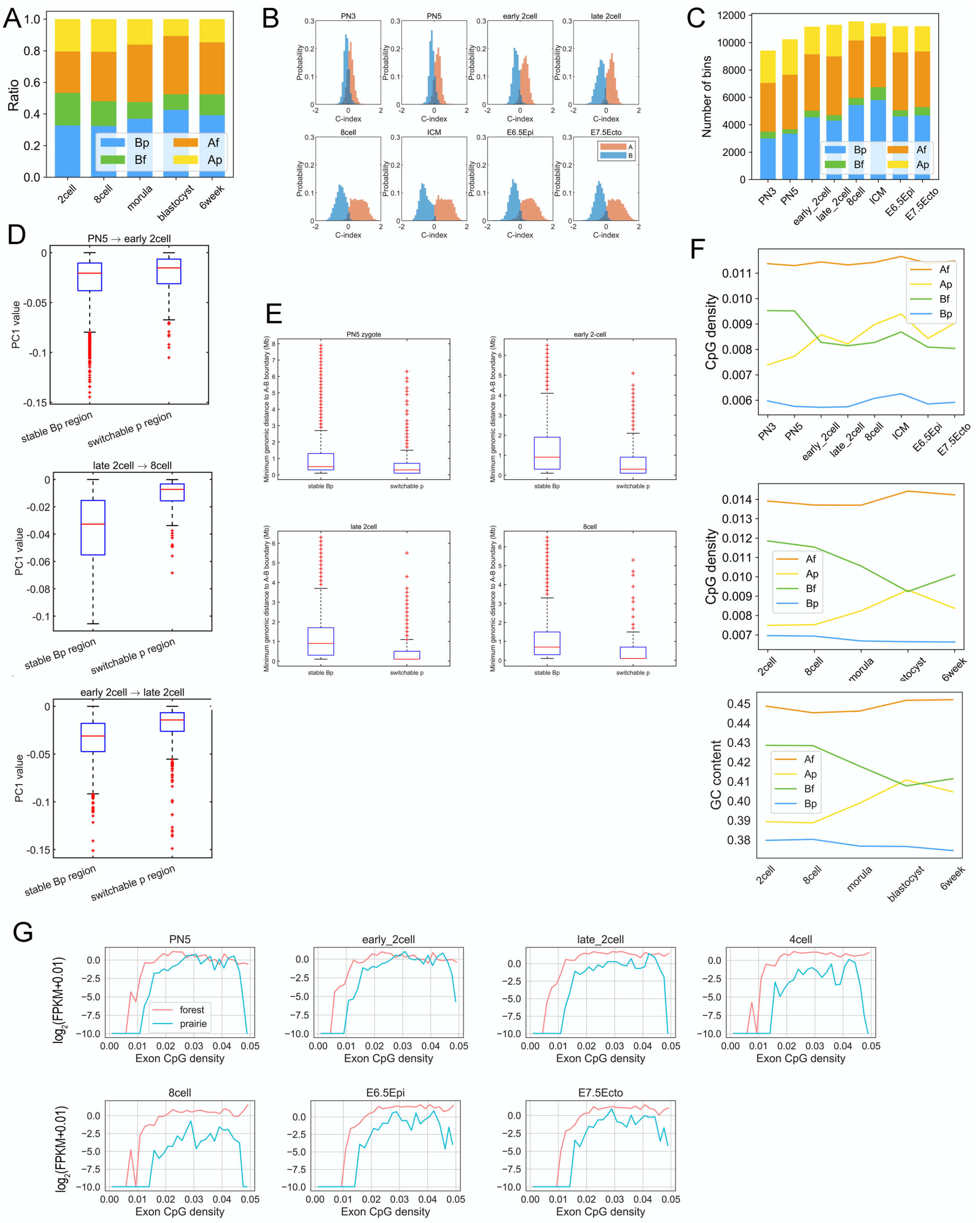

**Figure S3. Compartment changes relate with domain segregation.** Related to Figure 2.

(A) The proportion of four sequence components, Af, Ap, Bf and Bp (calculated according to the positioning of forests and prairies in compartments A and B) for each stage in human embryo development.

(B) The compartment index (C-index) distribution of compartment A and B regions during mouse embryo development.

(C) The number of bins belonging to sAf, sAp, sBf and sBp, in each stage during mouse embryo development.

(D) The PC1 value of stable Bp (i.e., prairie regions constantly remaining in compartment B in cell transition) and switchable p regions (i.e., prairie regions located in compartment B in early stage while switching to compartment A in late stage) in the early stage, during the transitions from PN5 to early 2-cell, from early 2-cell to late 2-cell and from late 2-cell to 8-cell.

(E) The minimum genomic distance between stable Bp region/switchable p region and A-B boundary. P-value = $3.6 \times 10^{-9}$, $1.6 \times 10^{-47}$, $7.2 \times 10^{-15}$ and $2.2 \times 10^{-7}$ for PN5, early 2-cell, late 2-cell and 8cell, respectively, by Welch's unequal variance test.

(F) The average CpG densities and GC contents of the four components in development. The four components, Af, Bf, Ap and Bp, are divided according to the positioning of forests and prairies in compartments A and B.

(G) The relation between gene expression level and gene exon CpG density.

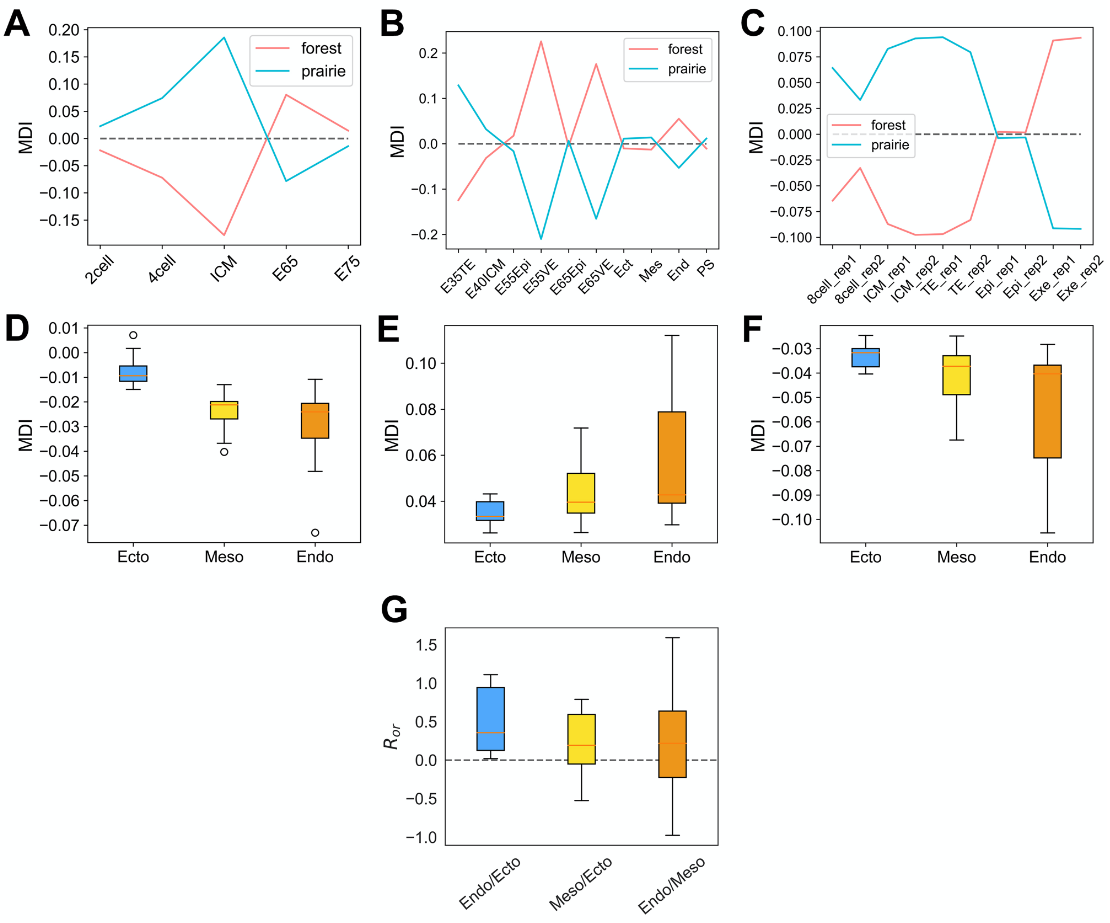


**Figure S4. Domain segregation and differentiated DNA methylation associated with germ layer formation.** Related to Figure 3.

(A, B, C) The average forest/prairie open sea methylation difference (MDI) at different stages during embryonic development. The methylation data are from three different groups: (A) Wang L et al., (B) Zhang Y et al., (C) Smith ZD et al (see Additional file 2: Data sources). The results all show that the absolute value of MDI increases from 2-cell to ICM stage, and then decreases from ICM to E6.5 epiblast stage.

(D, E, F) The box plots of average MDI for human and mouse differentiated cells. The cells considered originate from ectoderm, mesoderm and endoderm respectively, for human prairies (D), mouse forests (E) and mouse prairie domains (F). The absolute values of MDI increase in the order of ectoderm, mesoderm, endoderm.

(G) The box plots of overall relative segregation ratio $R_{or}$ of one tissue over another for prairies. The three categories are tissues derived from endoderm and mesoderm over those derived from ectoderm, and endoderm over mesoderm.

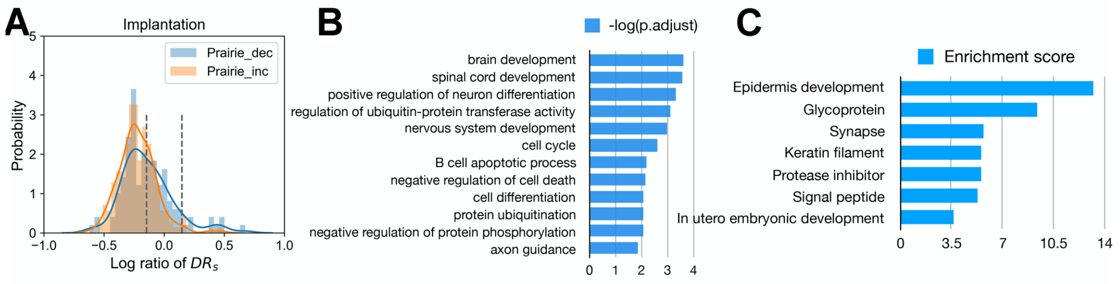


**Figure S5. Implantation-related domain mixing in differentiation.** Related to Figure 5.

(A) The domain segregation ratio changes of prairie domains harboring up- or down-regulated genes during implantation.

(B) The functional assignment of up-regulated genes in the $P_{mix}$ regions in implantation.

(C) The functional annotation clustering of genes in prairie domains moving from compartment B to A during implantation.

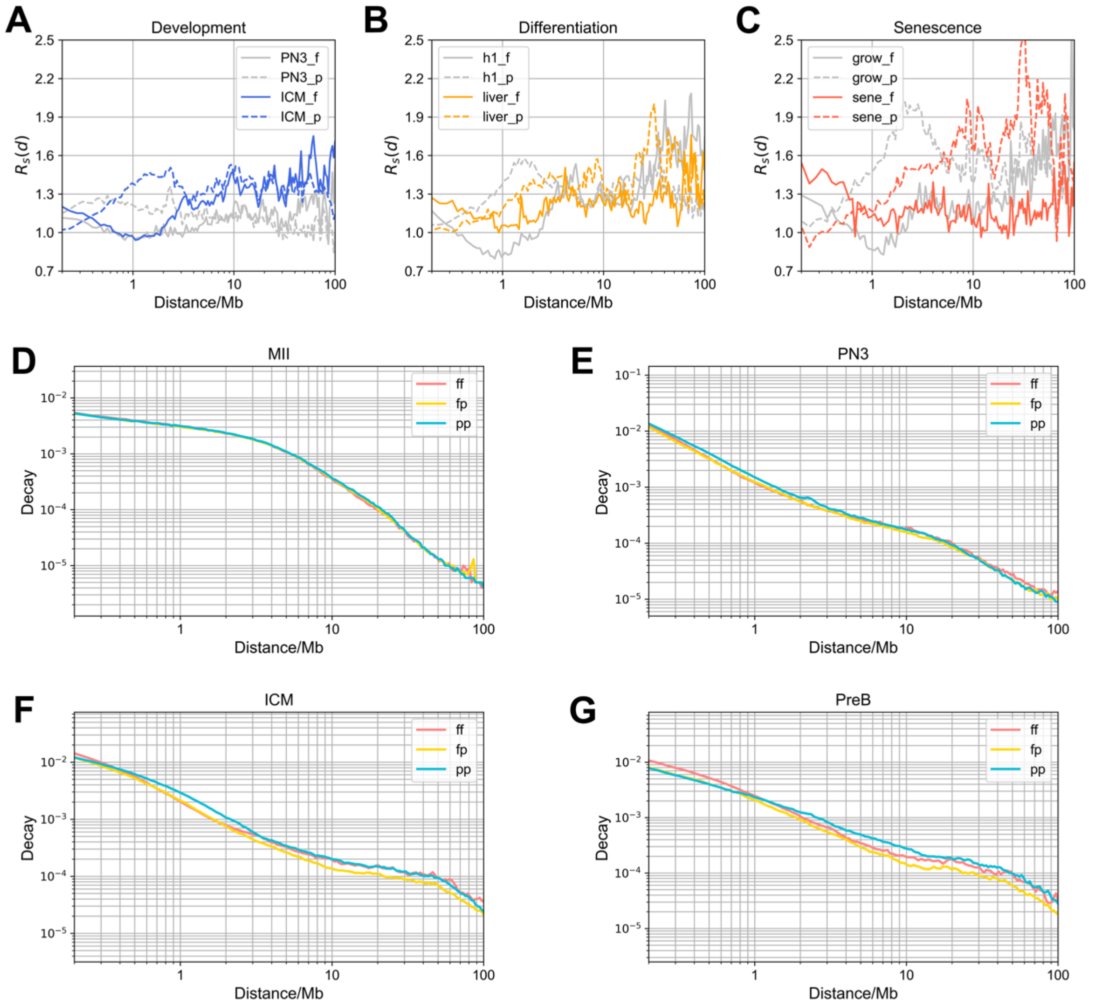


**Figure S6. Chromatin structural patterns at different stages.** Related to Figure 6.

(A-C) The distance-dependent segregation ratio changes during development(A), differentiation(B) and senescence(C). From zygote to ICM, chromatin is first seen to segregate clearly at 1~3Mb length scales. During differentiation, chromatin segregation occurs at long distances. In senescence, long-range segregation enhances further. Overall, a gradual establishment of long-range contacts occurs from birth to senescence.

(D-G) The decay of contacts between same-type genomic domains (ff and pp) and different-type genomic domains (fp) for MII (D), PN3 (E), ICM(F) and pre-B cell(G). After fertilization, contacts at short distances first increase. At ICM, long-range contacts have been gradually established at more than 10 Mb. Finally, in differentiation, contacts increase at genomic distances of 1~10 Mb.

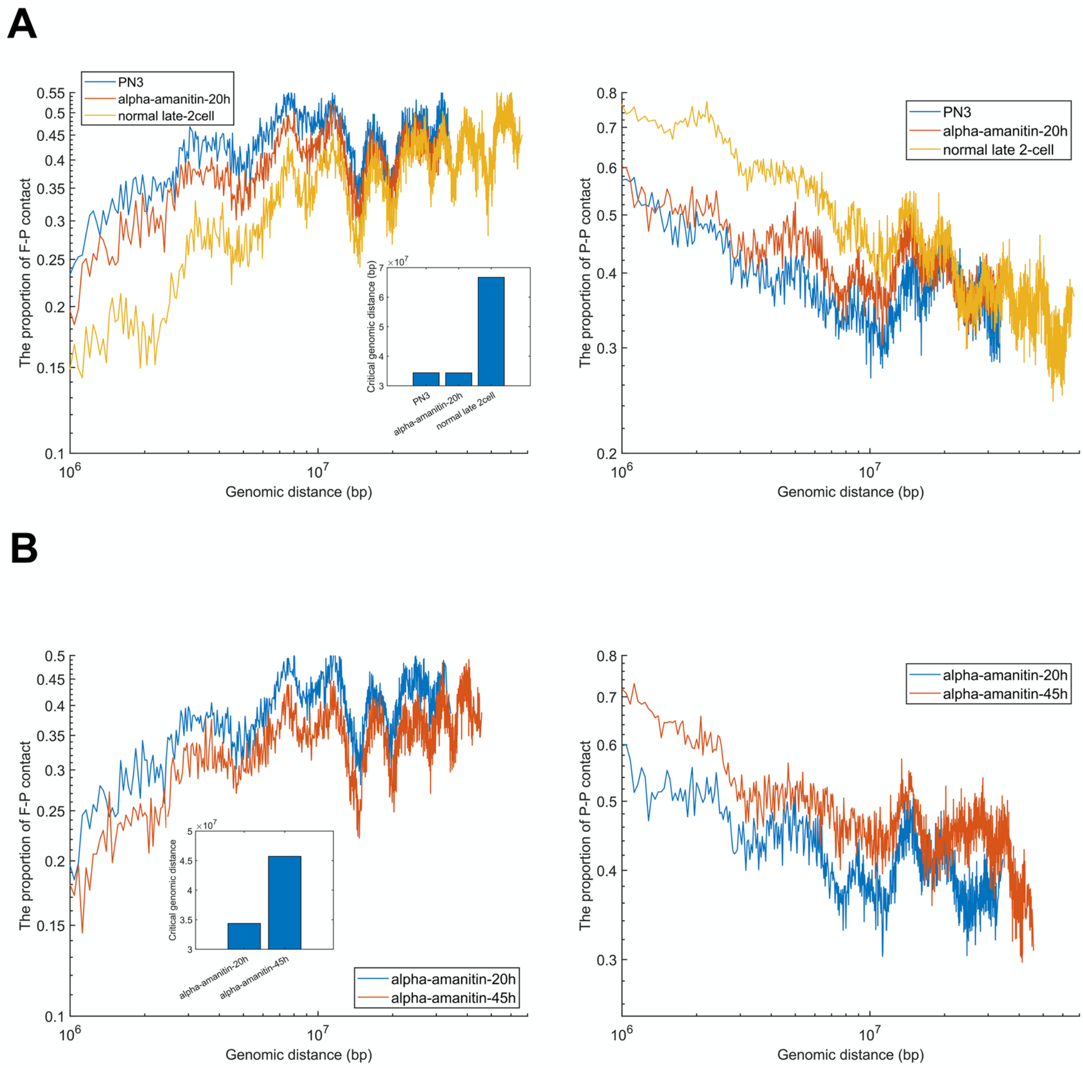


**Figure S7. The association between transcription inhibition and genome architecture.**

(A) The proportion of F-P and P-P spatial interactions within the top 10% contact probabilities under one certain genomic distance, at PN3, $\alpha$-amanitin-treated-20h cell and normal late 2-cell. Inner plot represents the critical genomic distance of PN3, $\alpha$-amanitin-treated-20h and normal late 2-cell.

(B) The proportion of F-P and P-P spatial interactions within the top 10% contact probabilities under one certain genomic distance at $\alpha$-amanitin-treated-20h cell and $\alpha$-amanitin-treated-45h cell. Inner plot represents the critical genomic distance of $\alpha$-amanitin-treated-20h and $\alpha$-amanitin-treated-45h.
